# Cellular diversity of the developing chick trigeminal ganglion at single-cell resolution

**DOI:** 10.64898/2026.02.01.702869

**Authors:** Arvind Arul Nambi Rajan, Erica J. Hutchins

**Author notes:** CORRESPONDENCE: Erica J. Hutchins.

## Abstract

**Background:** The trigeminal ganglion (TG) is a structure of the peripheral nervous system, composed of neuronal and non-neuronal cell types, that integrates sensory input from the face and jaw. The developing TG is derived from two embryonic cell populations: neural crest and cranial placode. Both populations play critical roles in TG development and must interact to coordinate changes in gene expression that regulate specification, differentiation, and organization. However, the molecular characteristics of the heterogeneous cell populations within the developing TG remain poorly defined.

**Results:** We performed single-cell RNA-sequencing (scRNA-seq) on TG from developing chick embryos at HH17. Our high-resolution dataset (14 clusters, ∼87000 cells) provides insight into cellular diversity within the developing TG. As expected, we identified placode-derived neurons as well as neural crest cells prior to neuronal differentiation. In addition to classic markers, we identified novel transcripts with unknown roles in TG development, including several long non-coding RNAs (lncRNAs).

**Conclusions:** We generated a single-cell atlas of the developing chick trigeminal ganglion during early axonogenesis and defined the transcriptomic states of its diverse cell populations. Our results provide a useful resource for better understanding the cell populations contributing to TG development and gene expression that drives cell identity and differentiation.

## 1. INTRODUCTION

The trigeminal ganglion (TG) is a vertebrate organ responsible for integrating sensory input (*e.g.*, touch, pain, temperature) from the face and relaying this information to the central nervous system^1–3^. The mature ganglia exist in pairs, with one TG per hemisphere of the head providing innervations to the face and jaw via three branches of nerves: ophthalmic (V1), maxillary (V2), and mandibular (V3)^4,5^. The TG is comprised of sensory neurons, which project axons through these three nerve branches, but also consists of non-neuronal cell types such as glia, immune cells, fibroblasts, and endothelial cells^6–8^. Dysfunction or activation of sensory neurons in the TG can lead to headache disorders which affect ∼40% of the population, as well as more rare conditions such as trigeminal neuralgia which also results in neuropathic pain^1,3^. Further, aberrant trigeminal gangliogenesis during development may also lead to deficits in pain or temperature sensation in the face^9^.

The trigeminal is just one of several cranial sensory ganglia^5^, which arise from two embryonic cell populations: neural crest and cranial placodes^10–16^. Neural crest are multipotent stem cells that are specified during neurulation but undergo tightly coordinated changes in gene expression to achieve epithelial-to-mesenchymal transition (EMT) and coordinate massive migrations that derive numerous terminal structures including TG^17–22^. Placodes are regions of thickened ectoderm that contribute to neurons of cranial sensory organs^10,12,13,23–25^. While some ganglia have neuronal contributions originating exclusively from either placode or neural crest, neurons of the epibranchial and trigeminal ganglia derive from both neural crest and placode progenitors^10,26,27^. In the case of these ganglia, placode differentiation into neurons precedes neural crest cell differentiation, which produces neurons and glia at a later stage^11,28,29^. In epibranchial ganglia, neurons derived from these distinct lineages are spatially segregated^10,11,26^; by contrast, neural crest- and placode-derived neurons are spatially intermingled within the TG^11,15,16^.

The interaction between placode and neural crest cells is complex and critical for proper development of the TG, and both progenitor populations are important determinants for condensation and morphogenesis of the organ^11,30,31^. The chick embryo has been an instrumental tool to understand neural crest and placode biology^10,13,14,16,21,24,26,30,31^, including elucidating time-resolved changes in gene expression during development of these embryonic cell populations^11,16,32–34^. Yet, the molecular gene expression within the diverse cell populations of the TG during this stage of development is incompletely understood.

Here, we present a high-resolution single-cell RNA-sequencing (scRNA-seq) atlas of the chick HH17 trigeminal ganglion, when placode-derived neurons are developing axons and neural crest cells have not yet begun to differentiate into neurons^13,28,31^. We believe this study will be a data-rich resource for other researchers in the field and hope it will facilitate future mechanistic and functional characterization of early trigeminal ganglion development.

## 2. RESULTS AND DISCUSSION

To investigate the cellular diversity of the developing trigeminal ganglion (TG), we performed single-cell RNA-sequencing (scRNA-seq) on dissected tissue from chick (*Gallus gallus*) embryos developed to stage HH17 (**Fig. 1A**). At this developmental stage, neural crest cells (SOX10^+^) and placode-derived cells (ISL1^+^) have condensed into a ganglion, with placode-derived neurons initiating axonogenesis (**Fig. 1B**) and further along in their differentiation trajectory than neural crest cells. As such, this is an ideal developmental time point to investigate the gene regulatory changes controlling the dynamics of these two progenitor populations during gangliogenesis. Two replicate sets of 20 pooled trigeminal ganglia were collected from left or right hemispheres of the embryo respectively. Only a single ganglion was dissected from each embryo, with left and right TG being collected separately for a total of four pooled samples (two left ganglia and two right ganglia samples). TG samples were then dissociated and processed for scRNA-seq using fixation and split-pool combinatorial barcoding from Parse Biosciences.

**Figure 1.**
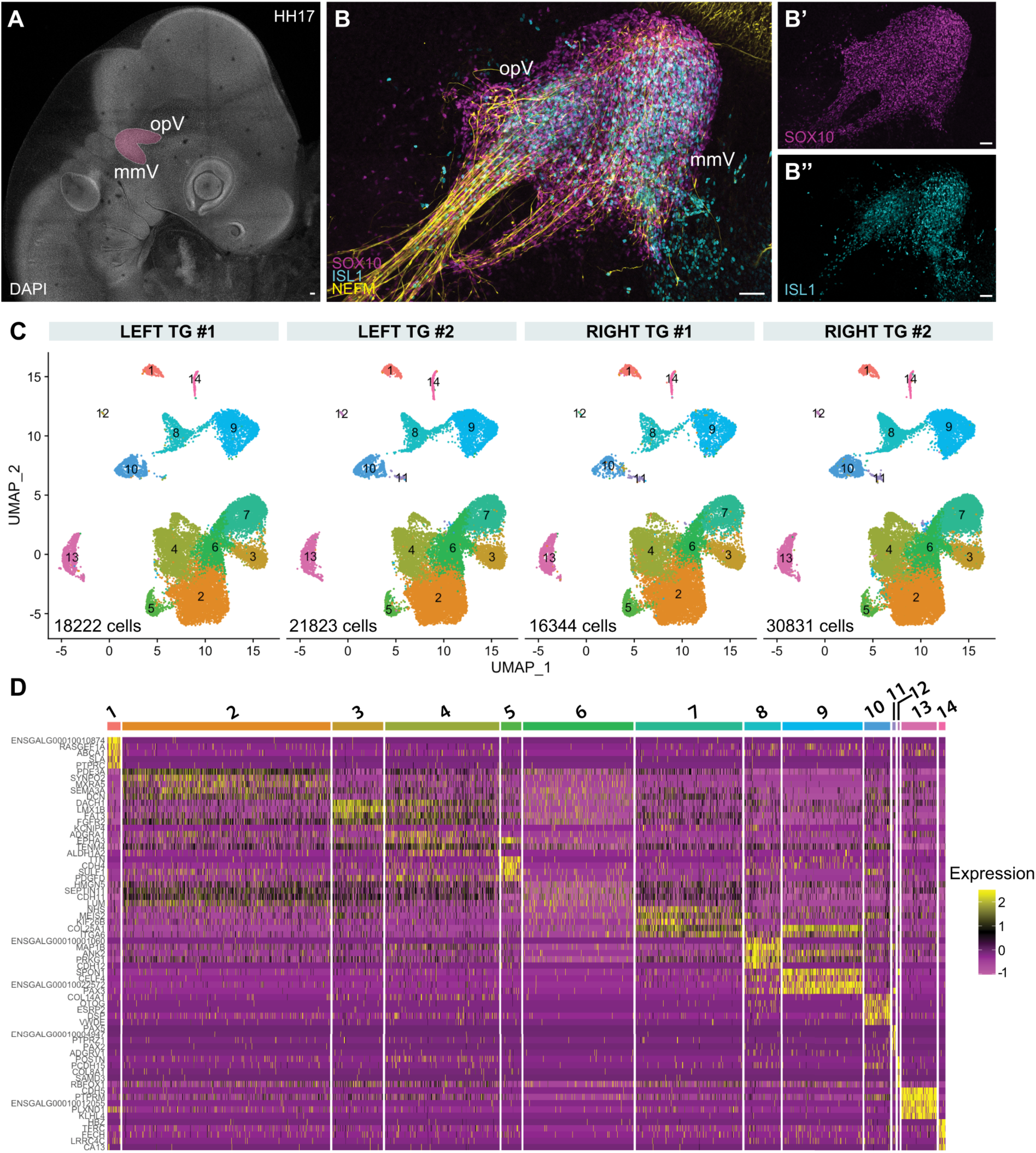
Single-cell RNA-sequencing of chick trigeminal ganglia identifies diverse cellular clusters. (**A**) Whole mount confocal maximum intensity projection of HH17 chick embryo stained with DAPI. Trigeminal ganglia (outlined and highlighted in magenta) were dissected for single-cell RNA-sequencing. (**B**) Whole mount confocal maximum intensity projection of a developing trigeminal ganglion at HH17, immunostained for SOX10 (**B’**; magenta), ISL1 (**B’’**; cyan), and NEFM (yellow). OpV, ophthalmic branch; MmV, maxillomandibular branch. Scale bar, 50 µm. (**C**) UMAP plots, split by sample, display 14 clusters identified in each sample. Total cell counts per sample are indicated on each plot. (**D**) Heatmap of top five differentially expressed marker genes across each cluster.

We recovered ∼87,220 total cells across the four samples (∼20,000-30,000 cells per sample) with 14 comparable clusters resolved for each sample after pre-processing and quality control (**Fig. 1C**; **Supplementary Fig. S1**). Cellular clusters were annotated using a combination of differential gene expression analysis (**Fig. 1D; Supplementary Table S1**) and marker gene identification based on previously published literature. Using this differential gene expression analysis and previously published literature as a guide, we generated a more curated custom annotation of cell types based on marker gene expression that represent diverse cellular populations, including: immune cells (*DOCK2*, *PTPRC*), fibroblasts (*COL1A1*, *LUM*, *DCN*), epithelial cells (*CLDN1*, *EPCAM*, *ESRP2*), endothelial cells (*FLT1*, *KDR*, *CDH5*), and erythrocytes (*HBBA*, *HBAD*, *HBA1*) (**Fig. 2A-B**)^6–8,16,35–43^.

**Figure 2.**
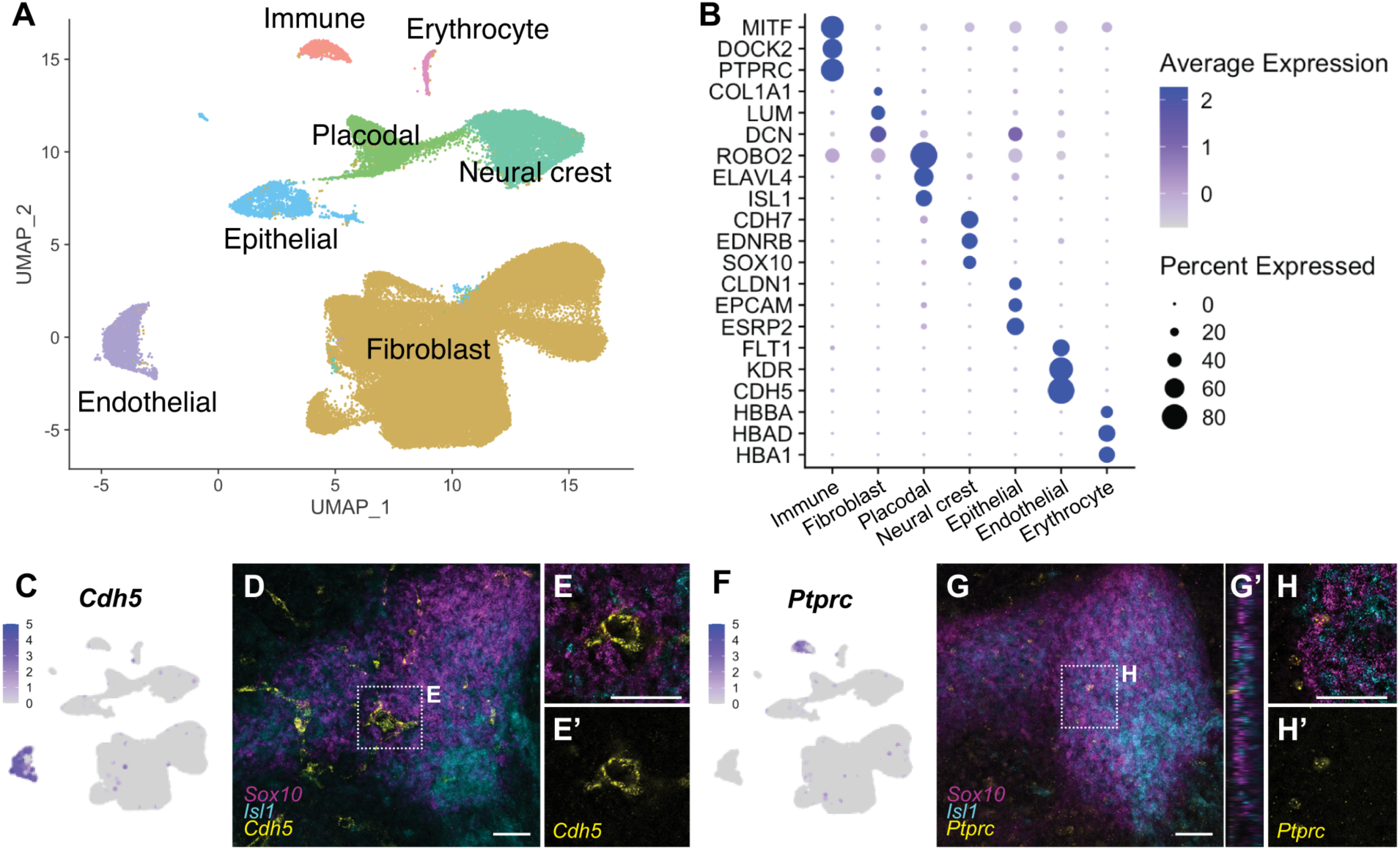
Cluster annotation and HCR validation for known marker genes. (**A**) UMAP representing combined data from all four single-cell RNA-seq samples, depicting manually-annotated clusters based on known marker gene expression. Seven major clusters were identified: immune cells, fibroblast cells, placodal cells, neural crest cells, epithelial cells, endothelial cells, and erythrocytes. (**B**) Dot plot depicting expression levels of marker genes for each manually-annotated cluster. (**C**) Feature plot of *Cdh5* expression, which is enriched in the endothelial cell cluster. (**D-E**) Whole mount confocal average intensity projection (**D**) and single Z plane (**E**) through (**D**) of a developing trigeminal ganglion at HH17, processed using HCR to visualize *Sox10* (magenta), *Isl1* (cyan), and *Cdh5* (yellow). Box in (**D**) indicates zoomed region in (**E**). (**F**) Feature plot of *Ptprc* expression, which is enriched in the immune cell cluster. (**G-H**) Whole mount confocal average intensity projection (**G**) and single Z plane (**H**) through (**G**) of a developing trigeminal ganglion at HH17, processed using HCR to visualize *Sox10* (magenta), *Isl1* (cyan), and *Ptprc* (yellow). Box in (**G**) indicates zoomed region in (**H**). Orthogonal projection in the YZ plane of (**G**) is displayed in (**G’**). Scale bar, 50 µm.

We validated a subset of these markers using hybridization chain reaction (HCR) to visualize their spatial expression *in vivo*. We observed *Cdh5^+^*endothelial cells within the developing TG (**Fig. 2C-E**). This is consistent with identification of endothelial cells in other scRNA-seq datasets from human and mouse TG^44^, as well as a requirement for endothelial cell interactions within other developing ganglia^45^. We also examined *Ptprc* expression to determine where immune cells localized at this stage of gangliogenesis. We found that immune cells were scattered around the developing TG with a few cells tightly adjacent to the surface of the ganglion, though none were detected within the ganglion interior (**Fig. 2F-H**), suggesting there may be neuro-immune crosstalk beginning at an early stage of trigeminal gangliogenesis.

Importantly, we further observed robust representation of neural crest and placode-derived cells expressing canonical markers (*e.g.*, *Sox10* and *Isl1*, respectively) (**Fig. 3A**). We detected placodal neurons based on expression of neurofilament *Nefm*, consistent with our immunostaining results (**Fig. 1B**), and pan-neuronal *Elavl4* within the *Isl1*^+^ cluster (**Fig. 3A**). HCR for *Fabp7* revealed that a subset of *Sox10*+ cells had developed into Schwann cell precursors (SCPs), marked by expression of *Fabp7* (**Fig. 3B-D**). The greater frequency of placode-derived neurons compared to neural crest-derived SCPs in our data set is consistent with the current understanding of temporal regulation of trigeminal ganglion development at this stage^31^. These data suggest that while placode-derived cells are further along in their developmental trajectories towards mature neurons compared to neural crest, a subset of neural crest are also beginning to differentiate into SCPs by HH17.

**Figure 3.**
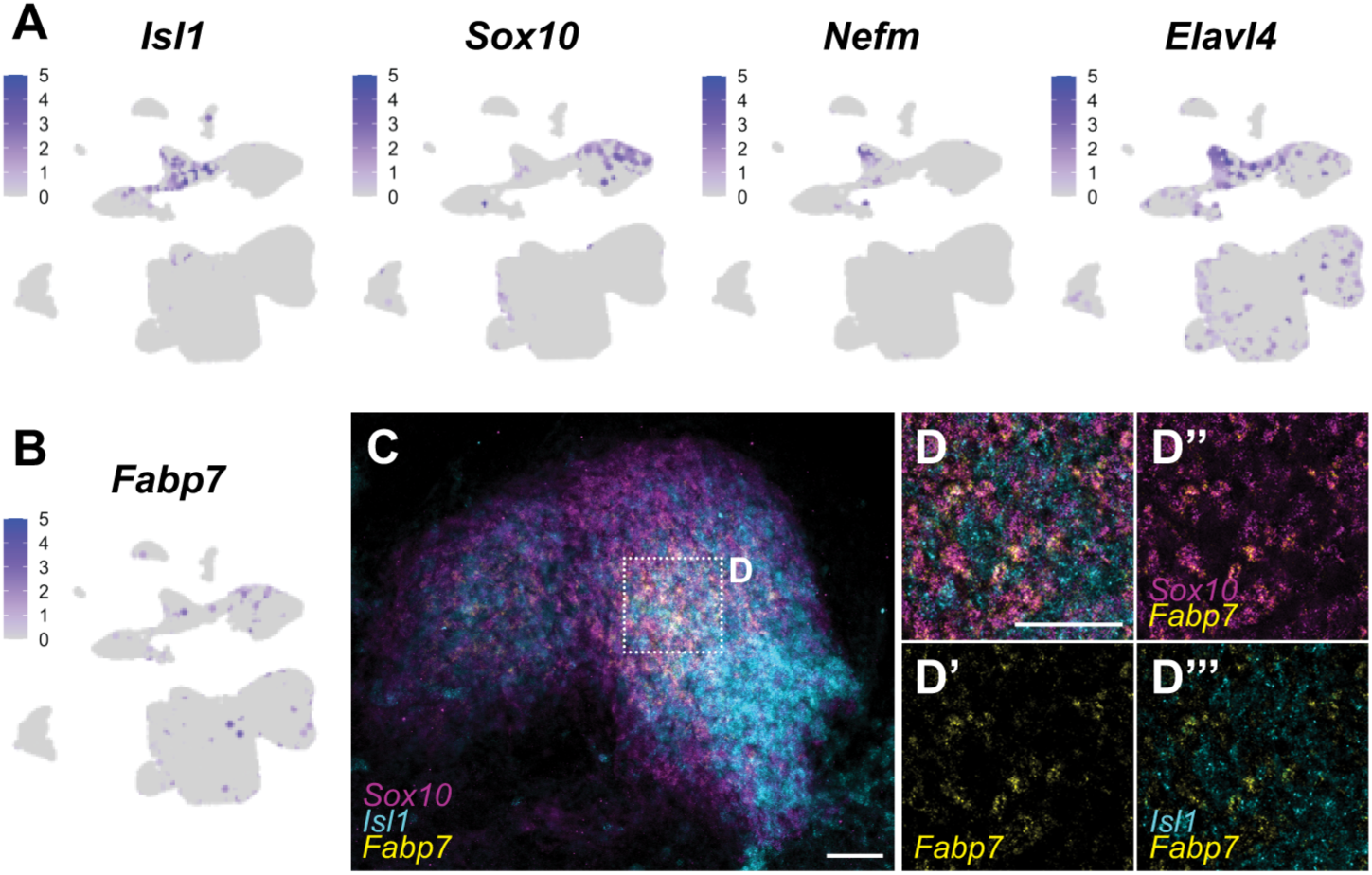
Feature plots and HCR validation for neural crest- and placode-derived marker genes. (**A**) Feature plots of *Isl1*, *Sox10*, *Nefm*, and *Elavl4* expression in HH17 trigeminal ganglion data. (**B**) Feature plot of *Fabp7* expression, an early marker for Schwann cell precursors. (**C-D**) Whole mount confocal average intensity projection (**C**) and single Z plane (**D**) through (**C**) of a developing trigeminal ganglion at HH17, processed using HCR to visualize *Sox10* (magenta), *Isl1* (cyan), and *Fabp7* (yellow). Box in (**C**) indicates zoomed region in (**D**). *Fabp7* expression overlaps with a subset of *Sox10*^+^ cells (**D’’**), but not *Isl1*^+^ cells (**D’’’**). Scale bar, 50 µm.

The robust detection of cellular populations that are historically challenging to isolate without significant manipulation or the need for single nuclei isolation (*e.g.*, neurons^46–48^) reflect the sensitivity and quality of the atlas presented in this study. Our data quality is further supported by the high degree of reproducibility across samples (**Supplementary Fig. S2**). It has been reported that asymmetric activity is observed in other neural crest-derived tissues (*e.g.*, the observed differences in left versus right cardiac neural crest ablation)^49,50^. Although we collected left and right ganglia from different embryos, we do not observe obvious differences between TG collected from opposite hemispheres of the head. Dot plots split by sample and cluster IDs reveal that similar percentage of cells in each cellular cluster express marker genes associated with that respective cell type (**Supplementary Fig. S2**). Though we are limited in the statistical power to confidently detect subtle changes in gene expression between left and right hemispheres (given our duplicate samples), we observe a strong correlation (Pearson coefficient R^2^ ≥ 0.97) between each condition and replicate (**Supplementary Fig. S2**). These data suggest that the differences in gene expression profiles between left and right ganglia are likely limited and that our samples each reflect highly reproducible results.

Though we broadly classify clusters 2-7 as fibroblasts during our cell type annotation, we observe neuronal and SCP markers (*e.g. Elavl4*, *Fabp7*) expressed in specific subclusters of this manually-curated classification. While we have focused this study on generation and general annotation of this resource, we encourage researchers in the field to further explore additional subpopulations within this dataset with the hope that this will be an information-rich resource to identify novel aspects of gene expression and molecular biology underlying gangliogenesis. To this end, and as a proof of principal of this concept, we performed further exploration of neural crest- and placode-derived populations within our atlas.

Given the tight correlation between all four samples sequenced, the samples were treated as four independent replicates in pseudobulk analysis to increase statistical power in comparing gene expression profiles of placode-derived vs neural crest-derived clusters (**Fig. 4A; Supplementary Table S2**). While many known neural crest specific factors were identified (*e.g.*, *CDH7*, *EDNRB*), other genes have no previously published role in neural crest biology and may reflect previously uncharacterized factors with biological functions in neural crest and the trigeminal ganglia (**Supplementary Table S2**). Of particular interest was the prevalence of long non-coding RNAs (lncRNAs) within our analysis, with some exhibiting very tight expression within neural crest- or placode-derived cell clusters. Upon inspection of three examples (*ENSGALG00010008935*, *ENSGALG00010014698*, and *ENSGALG00010024337*), we observed that these chick lncRNA loci are located in genomic regions syntenic with mammalian microRNA (miRNA) host genes and contain conserved intragenic miRNAs: *ENSGALG00010008935* hosts miR-99a, miR-let7c, and miR-125b-2; *ENSGALG00010014698* hosts miR-181a-1 and miR-181b-1; and *ENSGALG00010024337* hosts miR-137. As such, we identify these lncRNAs here as *MIR99AHG*, *MIR181A1HG*, and *MIR137HG*, respectively.

**Figure 4.**
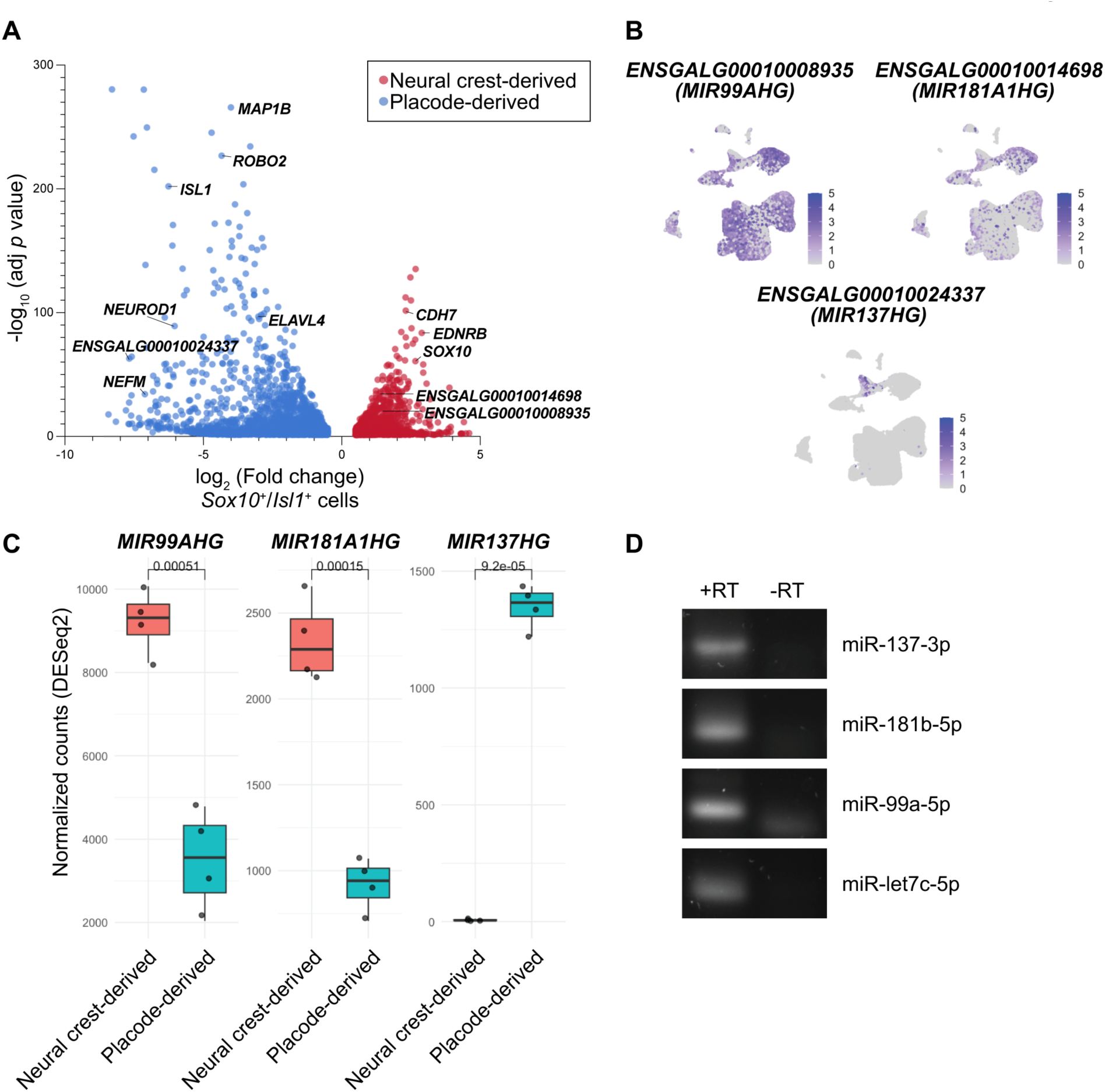
Differential expression analysis between pseudobulked neural crest and placode clusters revealed enriched lncRNA transcripts. (**A**) Volcano plot of differentially expressed genes between pseudobulked neural crest and placode clusters. 17253 genes are displayed after removing those that failed DESeq2 independent filtering (padj = NA). A selection of genes known to be enriched in neural crest- and placode-derived cells are indicated. Three novel long noncoding RNAs (lncRNAs) identified by differential expression analysis (*ENSGALG00010008935*, *ENSGALG00010014698*, and *ENSGALG00010024337*) are also indicated. (**B**) Feature plots of *ENSGALG00010008935*, *ENSGALG00010014698*, and *ENSGALG00010024337* expression in HH17 trigeminal ganglion data. Based on synteny, we identified these lncRNAs as *MIR99AHG*, *MIR181A1HG*, and *MIR137HG*, respectively. (**C**) Box plot depicting normalized counts for each lncRNA between pseudobulked neural crest and placode clusters. Individual data points represent each sample (*i.e.*, left TG#1, left TG #2, right TG #1, right TG #2). *P* values from *t*-test comparing neural crest and placode pseudobulked clusters are denoted, revealing statistically significant differences in expression for each of the lncRNAs. (**D**) Mature miRNA-specific RT-PCR (+RT) was performed on total RNA extracted from dissected trigeminal ganglia to confirm expression and processing of miRNAs hosted within each lncRNA. Negative (-RT) controls for each miRNA target were processed in parallel by omitting addition of reverse transcriptase.

Feature plots of *MIR99AHG* and *MIR181A1HG* show expression in (though not limited to) the neural crest-derived cluster, whereas *MIR137HG* shows tight expression largely limited to the placode-derived cluster of the single-cell atlas (**Fig. 4B**); given that miR-137 has been reported to be neuronally enriched^51^, we speculate *MIR137HG* expression in the trigeminal ganglion may be restricted to differentiated neurons. Comparisons of normalized counts for *MIR99AHG* and *MIR181A1HG* using DESeq2 revealed 2.8- and 2.6-fold higher expression, respectively, in pseudobulked neural crest-derived cells relative to placode-derived cells (*p* < 0.01); in contrast, *MIR137HG* had a 200-fold higher expression in placode-derived cells relative to neural crest (*p* < 0.001) (**Fig. 4C**).

To determine if the miRNAs hosted within these genes are expressed and processed into mature miRNAs during this stage of TG development, we performed RT-PCR on RNA extracted from dissected TG. Specifically targeting mature isoforms of candidate miRNAs nested within the aforementioned lncRNAs, we confirmed expression of miR-137-3p, miR181b-5p, miR-99a-5p, and miR-let7c-5p (**Fig. 4D**) in the developing trigeminal ganglion. This complements recent work which identified expression of miR181a-5p, miR-99a-5p, and miR-let7c-5p in neural crest-derived cells from small RNA sequencing at a similar stage of trigeminal ganglion development^52^. Together, these data raise the possibility for complex post-transcriptional mechanisms regulating neural crest and placode cell fate to drive trigeminal gangliogenesis. Future functional experiments will be critical in further defining the roles of these lncRNAs in the TG and validating this potentially novel post-transcriptional paradigm.

## 3. CONCLUSIONS

In this study, we present a high-resolution single-cell transcriptomic atlas of the HH17 trigeminal ganglion. Through analysis of diverse cellular populations defining early stages of trigeminal gangliogenesis, we identified potential post-transcriptional mechanisms that may be influencing cell fate decisions of these embryonic cells. Post-transcriptional regulators such as RNA-binding proteins and miRNAs have recently been implicated as important regulators of neural crest and trigeminal ganglion development^52–57^. Future studies will be needed to establish functional roles for novel proteins identified as marker genes in neural crest and placode clusters and elucidate the extent to which lncRNAs facilitate post-transcriptional control of these cell types. The dataset generated in this study contains novel information on additional cellular subpopulations within the trigeminal ganglion, which we think will be a rich resource for those studying neural crest or placode biology, gene expression, or gangliogenesis broadly. We encourage those in the field to explore the data generated here, and hope that it will facilitate continued investigation of trigeminal gangliogenesis.

## 4. EXPERIMENTAL PROCEDURES

### 4.1. Model Organism

Fertilized chicken eggs (*Gallus gallus*, Rhode Island Red breed) were purchased locally from Petaluma Farms (Petaluma, CA). Embryos were incubated in a humidified incubator set to 100°F until desired Hamburger-Hamilton (HH) stage^58^ was reached (∼66 hours).

### 4.2. Sample collection and single cell suspension preparation

Live embryos were excised from eggs at HH17 on Whatman filter paper as described^54,59,60^ and stored in Ringer’s solution until further processing. Pooled left and right trigeminal ganglia (TG) samples were generated by microdissection of 20 TG from separate embryos (left and right ganglia were never collected from the same embryo). Ganglia were collected and stored on ice in pre-chilled Ringer’s solution until two duplicate pooled left and right samples of 20 TG each were collected. Tissues were washed three times in pre-chilled DPBS and then enzymatically dissociated in 500µl 1X Accumax (Cat# AM105) supplemented with 25U RQ1 DNase (Cat# M6101) and 250U SUPERase-In (Cat# AM2694) at 37°C for 21 minutes with gentle pipette mixing every 5 minutes. Single cell suspensions were stored on ice while briefly manually counting cells using a hemocytometer and checking viability with trypan blue staining before proceeding to cell fixation for Parse Biosciences Split Pool Barcoding described below. Each 20 TG sample was confirmed to contain approximately 300,000 viable cells after dissociation and prior to fixation.

### 4.3. Parse single-cell fixation and RNA-sequencing

Single cell suspensions were processed with Parse Evercode Cell Fixation v3 kits per manufacturer’s instructions. All centrifugation steps were performed at 300xG in a swinging bucket rotor at 4°C and all cell straining steps were conducted using PluriSelect pluriStrainer mini with 20µm mesh size (Cat# 43-10020-40). Fixed samples were frozen in a Mr. Frosty Freezing Container at -80°C until sufficient sample numbers were generated to proceed with barcoding.

Fixed samples were thawed and processed for split pool barcoding using Parse Evercode WT 100K v3 kits per manufacturer’s protocols. To target a total of ∼100,000 barcoded cells output from the kit, we loaded approximately ∼87,000 fixed and viable cells per sample into round 1 of split pool barcoding (∼350,000 cells total across all 4 samples). Libraries were size selected using SPRIselect Reagent (Cat# B23317) when instructed. Library size distribution QC was conducted using Agilent Bioanalyzer High Sensitivity DNA Kit (Cat# UFUCOP-5067-4626) and quantified using Qubit dsDNA HS (High Sensitivity) Assay kit (Cat# Q33230). A total of 8 final sublibraries were generated and sequenced at a sequencing depth of 2 billion paired-end reads (20,000 reads/cell for 100,000 cells).

### 4.4. Transcriptomic Data Analysis

FASTQ files underwent barcode correction, read alignment, read deduplication, and transcript quantification using Parse Biosciences Trailmaker Pipeline Module (v1.6.2). The GRCg7b genome assembly was used as the reference genome for alignment. Additional data processing was done using the Parse Biosciences Trailmaker Insights Module. A cell size distribution filter using default settings was first applied to filter out cells with fewer than 331, 290, 365, and 374 transcripts in left TG sample #1, left TG sample #2, right TG sample #1, and right TG sample #2, respectively. These values were determined using barcode rank plots for each sample to identify cells with low transcript counts. Cells were next filtered based on the number of genes vs transcripts using the default third-order spline regression model where the prediction interval for defining outliers in the distribution was set to 0.999. A doublet filter using scDblFinder was then used to remove doublets with a probability threshold of 0.7. Resulting data was further processed in R (v4.4.0) with Seurat package (v4).

Data normalization, principal component analysis (PCA), and data integration was performed using Seurat and Harmony ^61,62^. A custom list of mitochondrial genes (*ND1, ND2, COX1, COX2, ATP8, ATP6, COX3, ND3, ND4L, ND4, ND5, ND6,* and *CYTB*) were used to estimate and plot percent.mt to confirm low mitochondrial content across all samples (<∼15%). LogNormalize was used to normalize, scale and log-transform feature expression measurements. 2000 highly variable genes were identified, and a linear transformation of the data was applied using the ScaleData() function. PCA was then performed on the scaled data, and the top 30 PCs were selected to define the dimensionality of the dataset. The top 30 PCs were then used in the FindNeighbors() function and cells were subsequently clustered using the Louvain algorithm by running FindClusters() in Seurat with a resolution of 0.3. Clusters were reordered according to their similarity using BuildClusterTree(). FindAllMarkers() was used to define differentially expressed genes in each cluster and identify marker genes. Top marker genes for each cluster were compared to published literature to manually annotate and merge Louvain clusters based on predicted cell type. Pseudobulk correlation analysis was used to assess reproducibility and variability across all four samples. For each sample, raw counts were aggregated across all cells and these aggregated counts were normalized using a log_1_(x+1) transformation to stabilize variance. Pairwise comparisons between samples were made by calculating Pearson correlation coefficients.

Differential gene expression analysis between placode-derived cells and neural crest-derived cells was also conducted using a pseudobulk approach to account for biological variation. Single-cell RNA-seq data was first subset to include only neural crest-derived and placode-derived clusters classified in **Fig. 2A** based on expression of known marker genes (including, but not limited to, *Sox10* and *Isl1*). Raw transcript counts were aggregated by biological replicate (chick) and cell type (collapsed) using AggregateExpression() function in Seurat. The resulting count matrix was analyzed employing the DESeq2 package^63^. To control for batch effects, the experimental design was modeled as ∼chick + collapsed, where chick represents the biological replicate and collapsed represents the cell population. The placode-derived group was defined as the reference level and differential expression was estimated using standard DESeq2 pipeline with results generated using the Wald test. Differential expression was considered significant if an adjusted *p* value (FDR) was < 0.05 and an absolute log_2_(fold change) was ≥ 0.5. For visualization of lncRNA expression profiles, normalized counts were extracted using counts() function within DESeq2. Box plots were generated using ggplot2 and ggpubr, with statistical significance determined using a two-sided t-test.

### 4.5. Immunohistochemistry (IHC) and Hybridization Chain Reaction (HCR)

Embryos were prepared for whole mount immunohistochemistry by fixation at room temperature for 20min with 4% paraformaldehyde in sodium phosphate buffer. Washes were performed in 1X TBST (0.5M Tris-HCl, 1.5M NaCl, 10mM CaCl2, 0.5% Triton X-100, 0.001% Thimerosal) and blocking was carried out in 10% donkey serum diluted in 1X TBST as described previously^54^. Primary antibodies were used to label NEFM (Thermo Cat# 13-0700; 1:200), SOX10 (Sigma Cat# HPA068898; 1:500), and ISL1 (DSHB Cat# 40.2D6, concentrate; 1:500). Species-specific secondary antibodies were labelled with Alexa Fluor 488 (1:1000), 568 (1:1000), or 647 (1:500) . Whole mount images were acquired on an inverted Zeiss LSM900 Airyscan 2. Images were adjusted for brightness/contrast and pseudocolored in FIJI^64^.

Hybridization Chain Reaction (HCR) v3.0 was performed per manufacturers’ protocol (Molecular Instruments) as described previously^54^. Prior to starting HCR, embryos were fixed in 4% paraformaldehyde at room temperature for 1 hour, then dehydrated by sequential washes in 25%, 50%, and 75% Methanol in PBST, and finally stored at least one overnight at -20°C. Embryos were treated with 10ug/mL proteinase K for 2 minutes and 30 seconds prior to post-fixation and downstream processing. Custom HCR v3.0 probes targeting *Ptprc*, *Fabp7*, and *Cdh5,* along with corresponding hairpins for fluorescent detection (488, 546 and 647), were designed by and ordered through Molecular Instruments^65^.

### 4.6. Total RNA extraction and mature miRNA-specific RT-PCR

Total RNA was extracted from 10 dissected and pooled trigeminal ganglia (5 left ganglia and 5 right ganglia, each from separate embryos, pooled together for one sample), collected in triplicate. TG were washed 3 times in 1X DPBS on ice followed by addition of 200µl RIPA buffer (50mM Tris-HCl, 150mM NaCl, 0.1% SDS, 1% Triton X-100) and tissue was disrupted on ice with plastic mortar and pestle. Phenol-chloroform extraction was used to isolate RNA. Mature miRNA-specific reverse transcription was performed per manufacturer’s instruction using Mir-X-miRNA First-Strand Synthesis kits (Cat# 638313). For PCR, the mRQ 3’ universal primers provided with the kits were used with 5’ miRNA target-specific primers. Primers for gga-miR-137-3p (5’-TATTGCTTAAGAATACGCGTAG), gga-miR-181b-1-5p (5’-AACATTCATTGCTGTCGGTGGG), gga-miR-99a-5p (5’-AACCCGTAGATCCGATCTTGTG), and gga-let7c-5p (5’-TGAGGTAGTAGGTTGTATGGTT) were designed to be the entire sequence of the mature target miRNA. Negative (-RT) controls for each miRNA target were processed in parallel by omitting addition of reverse transcriptase to control for non-specific amplification or genomic DNA contamination. PCR reactions were performed using Q5 Polymerase (Cat# cat. no. 638313).

## ACKNOWLEDGMENTS

The authors are supported by the National Institutes of Health (NIH) R00DE028592 (EJH).

## DATA AVAILABILITY STATEMENT

Single cell RNA-seq raw sequencing files have been deposited at NCBI Gene Expression Omnibus GSE318002.

## AUTHOR CONTRIBUTIONS

Conceptualization (AANR, EJH), Writing – Original Draft (AANR), Writing – Review & Editing (EJH), Investigation and Analysis (AANR), Funding (EJH).

## Supplementary Material

Contains two Supplementary Figures and two Supplementary Tables

### Supplemental Table legends

**Supplemental Table S1. Trigeminal ganglion scRNA-seq marker gene expression and cluster annotation.**

**Supplemental Table S2. Differentially expressed genes from pseudobulk analysis comparing neural crest- and placode-derived cells.**

**Supplemental Figure S1.**
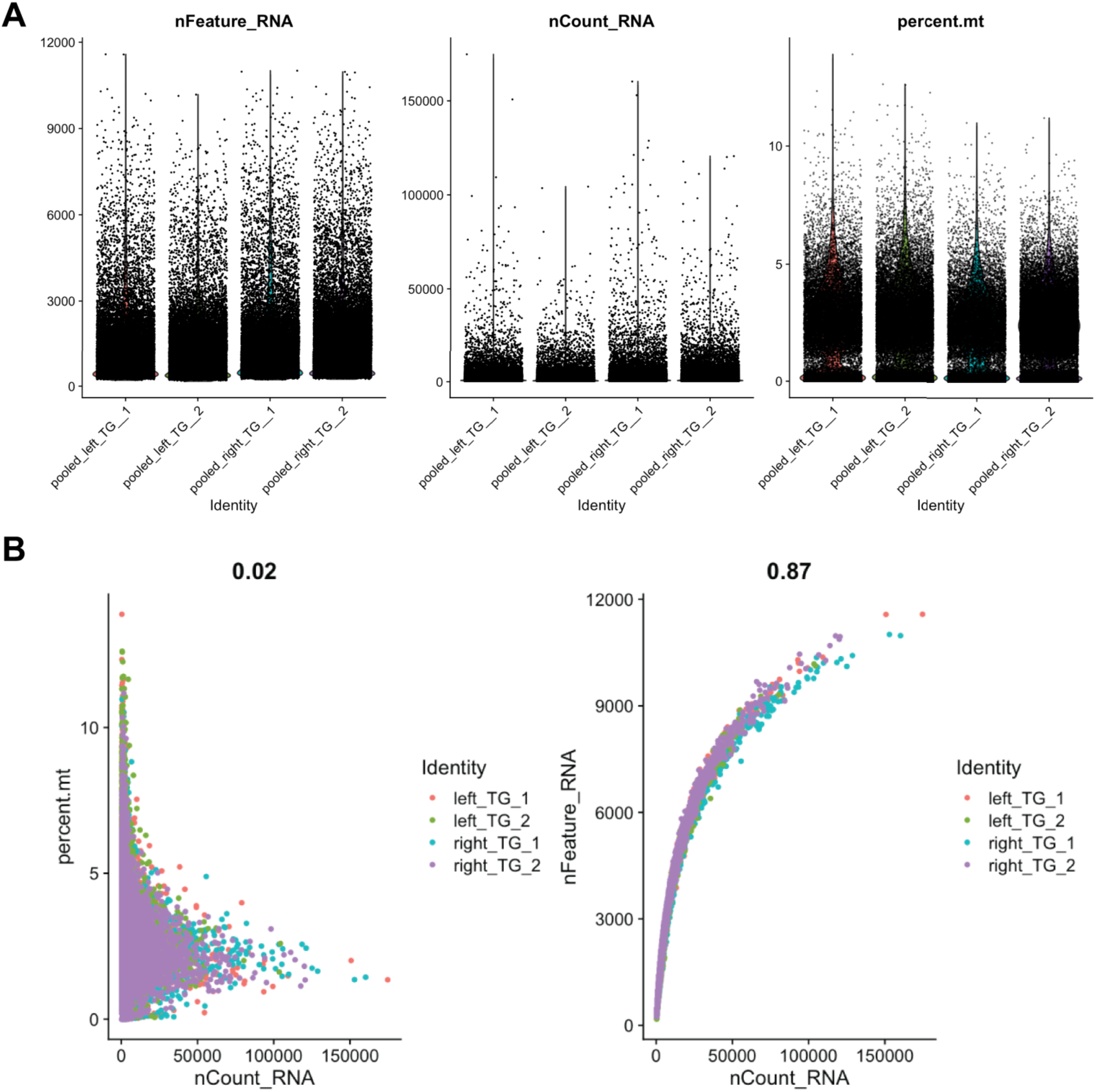
Quality Control (QC) of single-cell RNA-sequencing libraries. (**A**) nFeature_RNA, nCount_RNA, and percent.mt distributions for each sample are provided for transparency and to show the data is of high quality. The low percent mitochondrial content across all samples (<15%) suggests cells were healthy and not dying. Both nFeature_RNA and nCount_RNA distributions appear reflective of ranges that are expected for healthy cells. The high sensitivity of the Parse technology is reflected in the high gene and feature detection counts. (**B**) nCount_RNA and percent.mt show very poor correlation (Pearson correlation coefficient = 0.02), while nFeature_RNA and nCount_RNA show strong correlation (Pearson correlation coefficient = 0.87), as expected for a high-quality dataset.

**Supplemental Figure S2.**
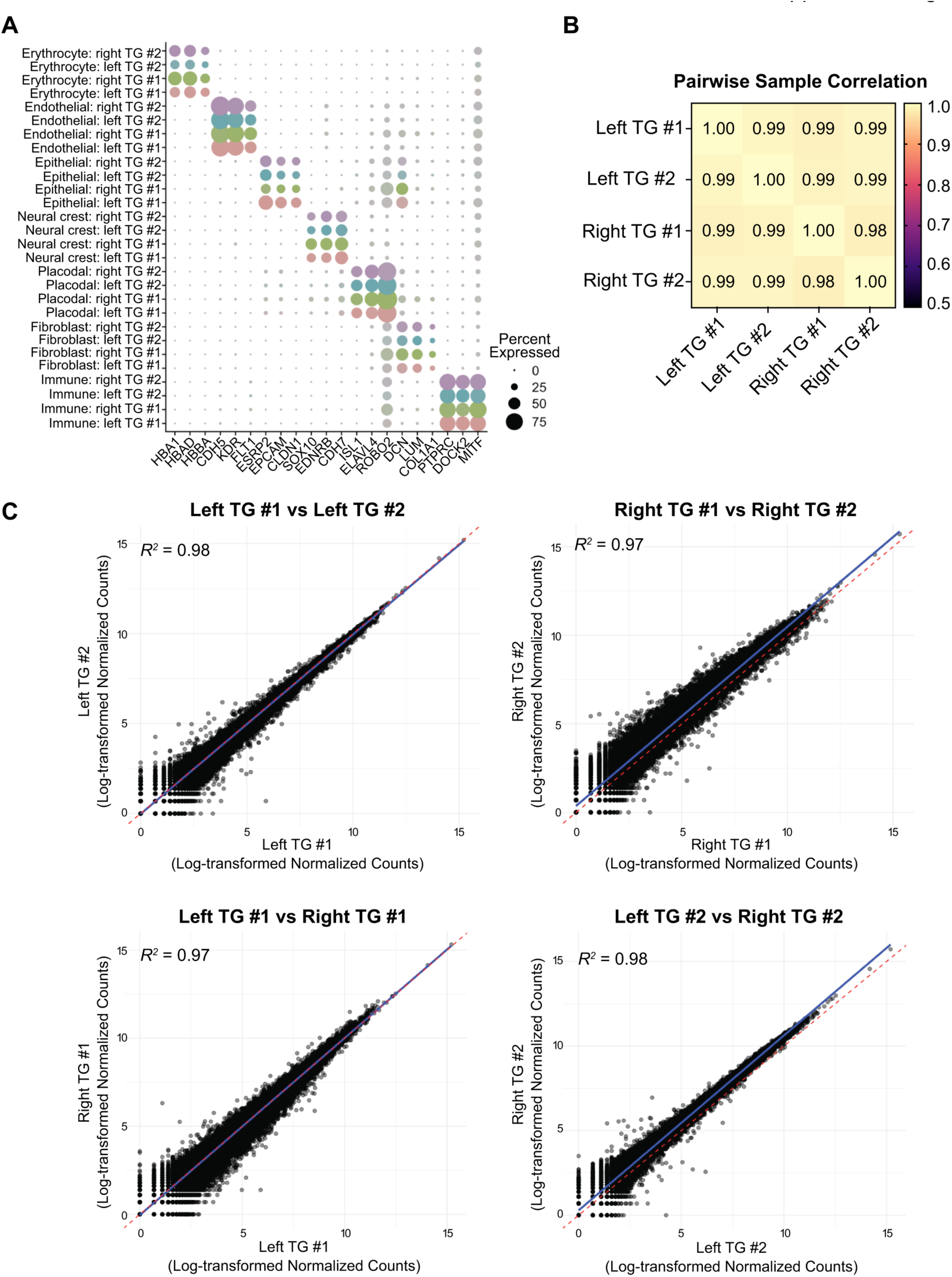
Left vs Right trigeminal ganglia (TG) and replicates all show tight correlation with each other. (**A**) Dot plot showing expression patterns of known marker genes split by sample and manually annotated cluster IDs. Similar patterns of marker gene expression in each cluster between left and right trigeminal ganglia (TG), as well as between replicates, are observed. (**B**) Heatmap of Pearson correlation on log-transformed pseudobulked data making pairwise comparisons between all samples sequenced. Left and right TG samples as well as all replicates show strong correlation and similarities between samples. (**C**) Representative scatterplots comparing log-transformed normalized counts for each gene between samples. Pearson correlation coefficients are denoted on each plot by the *R^2^* value.

## REFERENCES

1. Messlinger K, Russo AF. Current understanding of trigeminal ganglion structure and function in headache. Cephalalgia. Nov 2019;39(13):1661–1674. 10.1177/0333102418786261.

2. Goto T, Oh SB, Takeda M, et al. Recent advances in basic research on the trigeminal ganglion. J Physiol Sci. Sep 2016;66(5):381–6. 10.1007/s12576-016-0448-1.

3. Harriott AM, Gold MS. Contribution of primary afferent channels to neuropathic pain. Curr Pain Headache Rep. Jun 2009;13(3):197–207. 10.1007/s11916-009-0034-9.

4. Romano N, Federici M, Castaldi A. Imaging of cranial nerves: a pictorial overview. Insights Imaging. Mar 15 2019;10(1):33. 10.1186/s13244-019-0719-5.

5. Vermeiren S, Bellefroid EJ, Desiderio S. Vertebrate Sensory Ganglia: Common and Divergent Features of the Transcriptional Programs Generating Their Functional Specialization. Front Cell Dev Biol. 2020;8:587699. 10.3389/fcell.2020.587699.

6. Bhuiyan SA, Xu M, Yang L, et al. Harmonized cross-species cell atlases of trigeminal and dorsal root ganglia. Sci Adv. Jun 21 2024;10(25):eadj9173. 10.1126/sciadv.adj9173.

7. Mecklenburg J, Shein SA, Malmir M, et al. Transcriptional profiles of non-neuronal and immune cells in mouse trigeminal ganglia. Front Pain Res (Lausanne). 2023;4:1274811. 10.3389/fpain.2023.1274811.

8. Yang L, Xu M, Bhuiyan SA, et al. Human and mouse trigeminal ganglia cell atlas implicates multiple cell types in migraine. Neuron. Jun 1 2022;110(11):1806–1821 e8. 10.1016/j.neuron.2022.03.003.

9. Leonard CE, Quiros J, Lefcort F, Taneyhill LA. Loss of Elp1 disrupts trigeminal ganglion neurodevelopment in a model of familial dysautonomia. Elife. Jun 17 2022;11. 10.7554/eLife.71455.

10. Hamburger V. Experimental analysis of the dual origin of the trigeminal ganglion in the chick embryo. J Exp Zool. Nov 1961;148:91–123. 10.1002/jez.1401480202.

11. Steventon B, Mayor R, Streit A. Neural crest and placode interaction during the development of the cranial sensory system. Dev Biol. May 1 2014;389(1):28–38. 10.1016/j.ydbio.2014.01.021.

12. Barlow LA. Cranial nerve development: placodal neurons ride the crest. Curr Biol. Mar 5 2002;12(5):R171–3. 10.1016/s0960-9822(02)00734-0.

13. D’Amico-Martel A, Noden DM. Contributions of placodal and neural crest cells to avian cranial peripheral ganglia. Am J Anat. Apr 1983;166(4):445–68. 10.1002/aja.1001660406.

14. Narayanan CH, Narayanan Y. Neural crest and placodal contributions in the development of the glossopharyngeal-vagal complex in the chick. Anat Rec. Jan 1980;196(1):71–82. 10.1002/ar.1091960108.

15. York JR, Yuan T, McCauley DW. Evolutionary and Developmental Associations of Neural Crest and Placodes in the Vertebrate Head: Insights From Jawless Vertebrates. Front Physiol. 2020;11:986. 10.3389/fphys.2020.00986.

16. Leonard CE, McIntosh A, Sanyal J, Taneyhill LA. The transcriptional landscape of the developing chick trigeminal ganglion. Dev Biol. Apr 2025;520:108–116. 10.1016/j.ydbio.2024.12.013.

17. Prasad MS, Charney RM, Garcia-Castro MI. Specification and formation of the neural crest: Perspectives on lineage segregation. Genesis. Jan 2019;57(1):e23276. 10.1002/dvg.23276.

18. Martik ML, Bronner ME. Riding the crest to get a head: neural crest evolution in vertebrates. Nat Rev Neurosci. Oct 2021;22(10):616–626. 10.1038/s41583-021-00503-2.

19. Tang W, Bronner ME. Neural crest lineage analysis: from past to future trajectory. Development. Oct 23 2020;147(20). 10.1242/dev.193193.

20. Simoes-Costa M, Bronner ME. Establishing neural crest identity: a gene regulatory recipe. Development. Jan 15 2015;142(2):242–57. 10.1242/dev.105445.

21. Gandhi S, Bronner ME. Insights into neural crest development from studies of avian embryos. Int J Dev Biol. 2018;62(1-2-3):183-194. 10.1387/ijdb.180038sg.

22. Mendez-Maldonado K, Vega-Lopez GA, Aybar MJ, Velasco I. Neurogenesis From Neural Crest Cells: Molecular Mechanisms in the Formation of Cranial Nerves and Ganglia. Front Cell Dev Biol. 2020;8:635. 10.3389/fcell.2020.00635.

23. Schlosser G. Induction and specification of cranial placodes. Dev Biol. Jun 15 2006;294(2):303–51. 10.1016/j.ydbio.2006.03.009.

24. Jidigam VK, Gunhaga L. Development of cranial placodes: insights from studies in chick. Dev Growth Differ. Jan 2013;55(1):79–95. 10.1111/dgd.12027.

25. Breau MA, Schneider-Maunoury S. Cranial placodes: models for exploring the multi-facets of cell adhesion in epithelial rearrangement, collective migration and neuronal movements. Dev Biol. May 1 2015;401(1):25–36. 10.1016/j.ydbio.2014.12.012.

26. Lwigale PY. Embryonic origin of avian corneal sensory nerves. Dev Biol. Nov 15 2001;239(2):323–37. 10.1006/dbio.2001.0450.

27. Karpinski BA, Bryan CA, Paronett EM, et al. A cellular and molecular mosaic establishes growth and differentiation states for cranial sensory neurons. Dev Biol. Jul 15 2016;415(2):228–241. 10.1016/j.ydbio.2016.03.015.

28. d’Amico-Martel A, Noden DM. An autoradiographic analysis of the development of the chick trigeminal ganglion. J Embryol Exp Morphol. Feb 1980;55:167–82.

29. D’Amico-Martel A. Temporal patterns of neurogenesis in avian cranial sensory and autonomic ganglia. Am J Anat. Apr 1982;163(4):351–72. 10.1002/aja.1001630407.

30. Shiau CE, Bronner-Fraser M. N-cadherin acts in concert with Slit1-Robo2 signaling in regulating aggregation of placode-derived cranial sensory neurons. Development. Dec 2009;136(24):4155–64. 10.1242/dev.034355.

31. Halmi CA, Leonard CE, McIntosh AT, Taneyhill LA. N-cadherin facilitates trigeminal sensory neuron outgrowth and target tissue innervation. Development. May 1 2025;152(9). 10.1242/dev.204369.

32. Williams RM, Lukoseviciute M, Sauka-Spengler T, Bronner ME. Single-cell atlas of early chick development reveals gradual segregation of neural crest lineage from the neural plate border during neurulation. Elife. Jan 28 2022;11. 10.7554/eLife.74464.

33. Williams RM, Candido-Ferreira I, Repapi E, et al. Reconstruction of the Global Neural Crest Gene Regulatory Network In Vivo. Dev Cell. Oct 21 2019;51(2):255–276 e7. 10.1016/j.devcel.2019.10.003.

34. Kotov A, Seal S, Alkobtawi M, et al. A time-resolved single-cell roadmap of the logic driving anterior neural crest diversification from neural border to migration stages. Proc Natl Acad Sci U S A. May 7 2024;121(19):e2311685121. 10.1073/pnas.2311685121.

35. Ji L, Xu S, Luo H, Zeng F. Insights from DOCK2 in cell function and pathophysiology. Front Mol Biosci. 2022;9:997659. 10.3389/fmolb.2022.997659.

36. Jia S, Liu J, Chu Y, Liu Q, Mai L, Fan W. Single-cell RNA sequencing reveals distinct transcriptional features of the purinergic signaling in mouse trigeminal ganglion. Front Mol Neurosci. 2022;15:1038539. 10.3389/fnmol.2022.1038539.

37. Fishwick KJ, Neiderer TE, Jhingory S, Bronner ME, Taneyhill LA. The tight junction protein claudin-1 influences cranial neural crest cell emigration. Mech Dev. Sep-Dec 2012;129(9-12):275–83. 10.1016/j.mod.2012.06.006.

38. Litvinov SV, Velders MP, Bakker HA, Fleuren GJ, Warnaar SO. Ep-CAM: a human epithelial antigen is a homophilic cell-cell adhesion molecule. J Cell Biol. Apr 1994;125(2):437–46. 10.1083/jcb.125.2.437.

39. Warzecha CC, Sato TK, Nabet B, Hogenesch JB, Carstens RP. ESRP1 and ESRP2 are epithelial cell-type-specific regulators of FGFR2 splicing. Mol Cell. Mar 13 2009;33(5):591–601. 10.1016/j.molcel.2009.01.025.

40. Zamir L, Singh R, Nathan E, et al. Nkx2.5 marks angioblasts that contribute to hemogenic endothelium of the endocardium and dorsal aorta. Elife. Mar 8 2017;6. 10.7554/eLife.20994.

41. Dejana E, Vestweber D. The role of VE-cadherin in vascular morphogenesis and permeability control. Prog Mol Biol Transl Sci. 2013;116:119–44. 10.1016/B978-0-12-394311-8.00006-6.

42. Ziegler BL, Valtieri M, Porada GA, et al. KDR receptor: a key marker defining hematopoietic stem cells. Science. Sep 3 1999;285(5433):1553–8. 10.1126/science.285.5433.1553.

43. Kerkela E, Lahtela J, Larjo A, Impola U, Maenpaa L, Mattila P. Exploring Transcriptomic Landscapes in Red Blood Cells, in Their Extracellular Vesicles and on a Single-Cell Level. Int J Mol Sci. Oct 25 2022;23(21). 10.3390/ijms232112897.

44. Yang L, Xu M, Bhuiyan SA, et al. Human and mouse trigeminal ganglia cell atlas implicates multiple cell types in migraine. Neuron. 2022;110(11):1806–1821.e8. 10.1016/j.neuron.2022.03.003.

45. Mompéo B, Engele J, Spanel-Borowski K. Endothelial cell influence on dorsal root ganglion cell formation. J Neurocytol. Feb 2003;32(2):123–9. 10.1023/b:neur.0000005597.28053.2a.

46. Kim N, Kang H, Jo A, Yoo SA, Lee HO. Perspectives on single-nucleus RNA sequencing in different cell types and tissues. J Pathol Transl Med. Jan 2023;57(1):52–59. 10.4132/jptm.2022.12.19.

47. Lake BB, Ai R, Kaeser GE, et al. Neuronal subtypes and diversity revealed by single-nucleus RNA sequencing of the human brain. Science. Jun 24 2016;352(6293):1586–90. 10.1126/science.aaf1204.

48. Oh JM, An M, Son DS, et al. Comparison of cell type distribution between single-cell and single-nucleus RNA sequencing: enrichment of adherent cell types in single-nucleus RNA sequencing. Exp Mol Med. Dec 2022;54(12):2128–2134. 10.1038/s12276-022-00892-z.

49. Solovieva T, Bronner ME. Congenital heart defects differ following left versus right avian cardiac neural crest ablation. Dev Biol. Mar 2025;519:30–37. 10.1016/j.ydbio.2024.12.003.

50. Liu JA, Rao Y, Cheung MPL, et al. Asymmetric localization of DLC1 defines avian trunk neural crest polarity for directional delamination and migration. Nat Commun. Oct 30 2017;8(1):1185. 10.1038/s41467-017-01107-0.

51. Smrt RD, Szulwach KE, Pfeiffer RL, et al. MicroRNA miR-137 regulates neuronal maturation by targeting ubiquitin ligase mind bomb-1. Stem Cells. Jun 2010;28(6):1060–70. 10.1002/stem.431.

52. Marquez RB, Sanchez Vasquez E, Alonso AM, et al. Core microRNAs regulate neural crest delamination and condensation in the developing trigeminal ganglion. Proc Natl Acad Sci U S A. Dec 9 2025;122(49):e2517668122. 10.1073/pnas.2517668122.

53. Guzman-Espinoza M, Kim M, Ow C, Hutchins EJ. "Beyond transcription: How post-transcriptional mechanisms drive neural crest EMT". Genesis. Feb 2024;62(1):e23553. 10.1002/dvg.23553.

54. Hutchins EJ, Gandhi S, Chacon J, Piacentino M, Bronner ME. RNA-binding protein Elavl1/HuR is required for maintenance of cranial neural crest specification. Elife. Oct 3 2022;11. 10.7554/eLife.63600.

55. Guzman-Espinoza M, Vander Wende HM, Pacheco JL, Roldan AO, Hutchins EJ. Characterization of Pumilio gene expression during early neural crest development. Differentiation. Jul-Aug 2025;144:100883. 10.1016/j.diff.2025.100883.

56. Bernardi YE, Sanchez-Vasquez E, Marquez RB, et al. miR-203 secreted in extracellular vesicles mediates the communication between neural crest and placode cells required for trigeminal ganglia formation. PLoS Biol. Jul 2024;22(7):e3002074. 10.1371/journal.pbio.3002074.

57. Rajan AAN, Hutchins EJ. Post-transcriptional regulation as a conserved driver of neural crest and cancer-cell migration. Curr Opin Cell Biol. Aug 2024;89:102400. 10.1016/j.ceb.2024.102400.

58. Hamburger V, Hamilton HL. A series of normal stages in the development of the chick embryo. J Morphol. Jan 1951;88(1):49–92.

59. Hutchins EJ, Bronner ME. Draxin alters laminin organization during basement membrane remodeling to control cranial neural crest EMT. Dev Biol. Feb 15 2019;446(2):151–158. 10.1016/j.ydbio.2018.12.021.

60. Hutchins EJ, Bronner ME. Draxin acts as a molecular rheostat of canonical Wnt signaling to control cranial neural crest EMT. J Cell Biol. Oct 1 2018;217(10):3683–3697. 10.1083/jcb.201709149.

61. Satija R, Farrell JA, Gennert D, Schier AF, Regev A. Spatial reconstruction of single-cell gene expression data. Nat Biotechnol. May 2015;33(5):495–502. 10.1038/nbt.3192.

62. Korsunsky I, Millard N, Fan J, et al. Fast, sensitive and accurate integration of single-cell data with Harmony. Nat Methods. Dec 2019;16(12):1289–1296. 10.1038/s41592-019-0619-0.

63. Love MI, Huber W, Anders S. Moderated estimation of fold change and dispersion for RNA-seq data with DESeq2. Genome Biol. 2014;15(12):550. 10.1186/s13059-014-0550-8.

64. 64. Schindelin J, Arganda-Carreras I, Frise E, et al. Fiji: an open-source platform for biological-image analysis. Nat Methods. Jun 28 2012;9(7):676–82. 10.1038/nmeth.2019.

65. Choi HMT, Schwarzkopf M, Fornace ME, et al. Third-generation in situ hybridization chain reaction: multiplexed, quantitative, sensitive, versatile, robust. Development. Jun 26 2018;145(12). 10.1242/dev.165753.

